# Probing the origin of matching functional jaws: roles of *Dlx5/6* in cranial neural crest cells

**DOI:** 10.1101/296665

**Authors:** Miki Shimizu, Nicolas Narboux-Nême, Yorick Gitton, Camille de Lombares, Anastasia Fontaine, Gladys Alfama, Taro Kitazawa, Yumiko Kawamura, Eglantine Heude, Lindsey Marshall, Hiroki Higashiyama, Youichiro Wada, Yukiko Kurihara, Hiroki Kurihara, Giovanni Levŕ

## Abstract

Gnathostome jaws derive from the first pharyngeal arch (PA1), a complex structure constituted by Neural Crest Cells (NCCs), mesodermal, ectodermal and endodermal cells. Here, to determine the regionalized morphogenetic impact of *Dlx5/6* expression, we specifically target their inactivation or overexpression to NCCs. NCC-specific *Dlx5/6* inactivation (*NCC*^Δ*Dlx5/6*^) generates severely hypomorphic lower jaws that present typical maxillary traits. Therefore, differently from the symmetric jaws obtained after constitutive *Dlx5/6* inactivation, *NCC*^Δ*Dlx5/6*^ embryos present a strikingly asymmetric mouth. Reciprocally, forced *Dlx5* expression in maxillary NCCs provokes the appearance of distinct mandibular characters in the upper jaw. We conclude that: 1) *Dlx5/6* activation in NCCs invariably determines lower jaw identity; 2) the morphogenetic processes that generate functional matching jaws depend on the harmonization of *Dlx5/6* expression in NCCs and in distinct ectodermal territories. The co-evolution of synergistic opposing jaws requires the coordination of distinct regulatory pathways involving the same transcription factors in distant embryonic territories.

## INTRODUCTION

The gnathostome skull is characterized by the presence of articulated, muscularized asymmetric jaws, capable to support predatory behaviours and feeding functions such as mastication and swallowing (Compagnucci, et al., 2013; Depew, et al., 2005). During development, upper and lower jaws derive from the maxillary and mandibular processes of the first pharyngeal arch (PA1) that give rise to the palatoquadrate dorsally and the Meckelian cartilage (MC) ventrally. Hox-free Neural Crest Cells (‘NCCs’) emigrating from the mesencephalic and anterior rhombencephalic neural folds (Ealba, et al., 2015; Fish, et al., 2014; Minoux and Rijli, 2010; Depew and Olsson, 2008; Noden and Trainor, 2005; Creuzet, et al., 2002; Kontges and Lumsden, 1996; Trainor and Tam, 1995; Couly, et al., 1993) colonize the PA1 and give rise to most facial cartilages, bones and tendons. Before migration, NCCs lack the topographic information needed to unfold jaw morphogenetic processes (Couly, et al., 2002). In the course of migration and after reaching their final destination in the craniofacial buds, NCCs contact mesodermal, ectodermal and endodermal cells. Morphogenetic cues exchanged between these different cellular populations result in the formation of functional and morphologically different upper and lower jaws capable to operate in synergy to assure feeding (Barske, et al., 2018; Sato, et al., 2008b; Tucker and Lumsden, 2004).

Surgical deletion and grafting of different regions of the embryonic anterior foregut endoderm and ectoderm have demonstrated that these epithelia convey to NCCs the topographic cues needed for PAs patterning and facial morphogenesis (Noden and Trainor, 2005; Ruhin, et al., 2003; Couly, et al., 2002). The molecular nature of these instructive signals is not completely elucidated, but experimental evidence suggests the involvement of bone morphogenetic proteins (BMPs), FGFs, endothelin-1 (Edn1), Notch/Delta and Sonic hedgehog (Depew and Simpson, 2006) signalling molecules.

In the mouse, loss of Edn1 and/or endothelin receptor type-A (Ednra) (Clouthier, et al., 1998; Kurihara, et al., 1994) result in the transformation of the lower jaw into a structure presenting major morphological hallmarks of an upper jaw such as absence of MC, zygomatic arch-like structures and vibrissae. These observations indicate that Edn1 signalling plays a major role in the specification of mandibular identity (Ozeki, et al., 2004; Ruest, et al., 2004). Remarkably, however, the upper jaw-like structures deriving from the mouse mandibular arch after Edn1-pathway inactivation are hypomorphic and clearly different from the normal upper jaw (Ruest, et al., 2004; Clouthier, et al., 1998). A further demonstration of the importance of Edn1-signalling for the establishment of jaw patterning comes from the constitutive activation of Ednra in upper jaw NCCs where its ligand, Edn1, is not normally present (Ozeki, et al., 2004). We reported that ectopic Ednra activation induces the transformation of maxillary into mandibular structures with duplicated MCs and dermatocranial jaws constituted by four, opposing dentary-like bones. In the same study we obtained a similar transformation forcing the expression of *Hand2*, a downstream target of the Edn1 pathway, in the Ednra-positive domain (Sato, et al., 2008a). It appears that Edn1 drives lower jaw development through a calmodulin-CamKII-histone deacetylase cascade that de-represses Mef2c, which then transactivates *Dlx5/6* expression (Hu, et al., 2015; Verzi, et al., 2007).

*Dlx* homeobox genes are vertebrate homologues of Drosophila *distal-less* which play a fundamental role in specifying dorsoventral PA1 patterning (Depew, et al., 2005; Merlo, et al., 2000). The six gnathostome *Dlx* genes are organized in bigenic tandems and are all expressed in NCCs of craniofacial primordia in spatially and temporally restricted patterns. *Dlx1* and *Dlx2* are present in both maxillary and mandibular NCCs while *Dlx5* and *Dlx6* are expressed only in mandibular NCCs. The targeted simultaneous inactivation of *Dlx5* and *Dlx6* (Beverdam, et al., 2002; Depew, et al., 2002) in all cells of the developing embryo, including NCCs and epithelial cells, leads to the transformation of the lower jaw into an upper jaw-like structure which, at variance with that obtained after *Edn1* or *Ednra* inactivation, presents a similar size and a remarkable symmetry compared to the existing upper jaw.

The transformations in jaw identity induced by the inactivation of Edn1 signalling are accompanied by the down regulation of *Dlx5* and *Dlx6* in NCCs (Fukuhara, et al., 2004; Ozeki, et al., 2004; Ruest, et al., 2004; Clouthier, et al., 1998); however, a territory of Edn-1 independent *Dlx5/6* expression is consistently maintained in the proximal PA1 (Heude, et al., 2010; Ozeki, et al., 2004). *Edn1* is expressed in the epithelium and mesodermal core of the mandibular part of PA1, whereas *Ednra* is almost exclusively expressed by NCCs suggesting that altered Edn1 signalling affects *Dlx5* and *Dlx6* expression only in NCCs and not in other cell types. Given the difference between the lower jaw morphology obtained in *Edn1/Ednra*mutants and *Dlx5/6* mutants, these findings suggest that the Edn1-dependent activation of *Dlx5/6* in NCCs is necessary to specify mandibular identity, but is insufficient to generate a normal lower jaw morphology (Sato, et al., 2008b; Fukuhara, et al., 2004; Ozeki, et al., 2004; Ruest, et al., 2004; Clouthier, et al., 1998). Interestingly it has been observed that, after inactivation of *Dlx5* and *Dlx6*, maxillary components are also affected despite the fact that these genes are not expressed by maxillary NCCs (Beverdam, et al., 2002; Depew, et al., 2002). This observation could be accounted for by the presence of signalling centres at the extremities of both the mandibular and maxillary arches; the so-called “Hinge and Caps” model of jaw organization (Compagnucci, et al., 2013; Fish, et al., 2011; Depew and Compagnucci, 2008). This model predicts the presence of two opposing morphogen gradients, one emanating from the region of the upper/lower jaw articulation (hinge) and one from the distal extremities of PA1 (caps); the origin and nature of these signals remain still elusive. By lineage analysis, we have shown that the maxillary arch epithelium harbours a cellular contingent derived from frontonasal *Dlx5*-expressing progenitors suggesting that transient *Dlx5/6* expression could program these epithelial cells to provide the cues needed for maxillary arch morphogenesis (Gitton, et al., 2014).

Here, we address the morphogenetic role of *Dlx5/6* expression in NCCs. To this end we first invalidate *Dlx5* and *Dlx6* specifically in NCCs and then we force ectopic expression of *Dlx5* in maxillary NCCs that do not normally express this gene. Our findings confirm that *Dlx5/6* expression in NCCs is necessary to specify mandibular identity and has an organizer role for craniofacial myogenesis. However, NCC-limited *Dlx5/6* expression is insufficient to determine normal jaw morphogenesis; coordination of *Dlx5/6* expression in NCCs and in other cellular components is essential for the development and evolution of a functional mouth.

## RESULTS

### Generation of mouse models with deregulation of *Dlx5/6* expression in NCCs

To induce ectopic expression of *Dlx5* in the NCC lineage, we crossed *ROSA^CAG-flox-Dlx5/+^* mice, which express *Dlx5* in a *Cre*-dependent manner, with *Wnt1^Cre^* mice (Chai, et al., 2000) to obtain the *NCC^Dlx5^* mouse line in which *Dlx5* should be expressed in premigratory neural crest cells (Fig. 1A). The expression of *Dlx5* in the pharyngeal arches was analyzed by quantitative RT-PCR (Fig. 1B) and whole-mount *in situ* hybridization (Fig. 1C, C’). RT-PCR was performed on dissected maxillary, mandibular and second pharyngeal (PA2) arches from E10.5 control and *NCC^Dlx5^* embryos. In all *NCC^Dlx5^* samples, *Dlx5* expression was higher than that found in the corresponding controls (Fig. 1B). By *in situ* hybridization, ectopic *Dlx5* expression was detected in the presumptive maxillary and nasal regions, well beyond the normal mandibular territory of expression (Fig. 1C, C’). This pattern is consistent with the distribution of *lacZ* expression in *Wnt-1::Cre/R26R* mice (Jiang, et al., 2000) suggesting that indeed *Dlx5* expression has been activated and maintained ectopically in NCCs.

**Figure 1:**
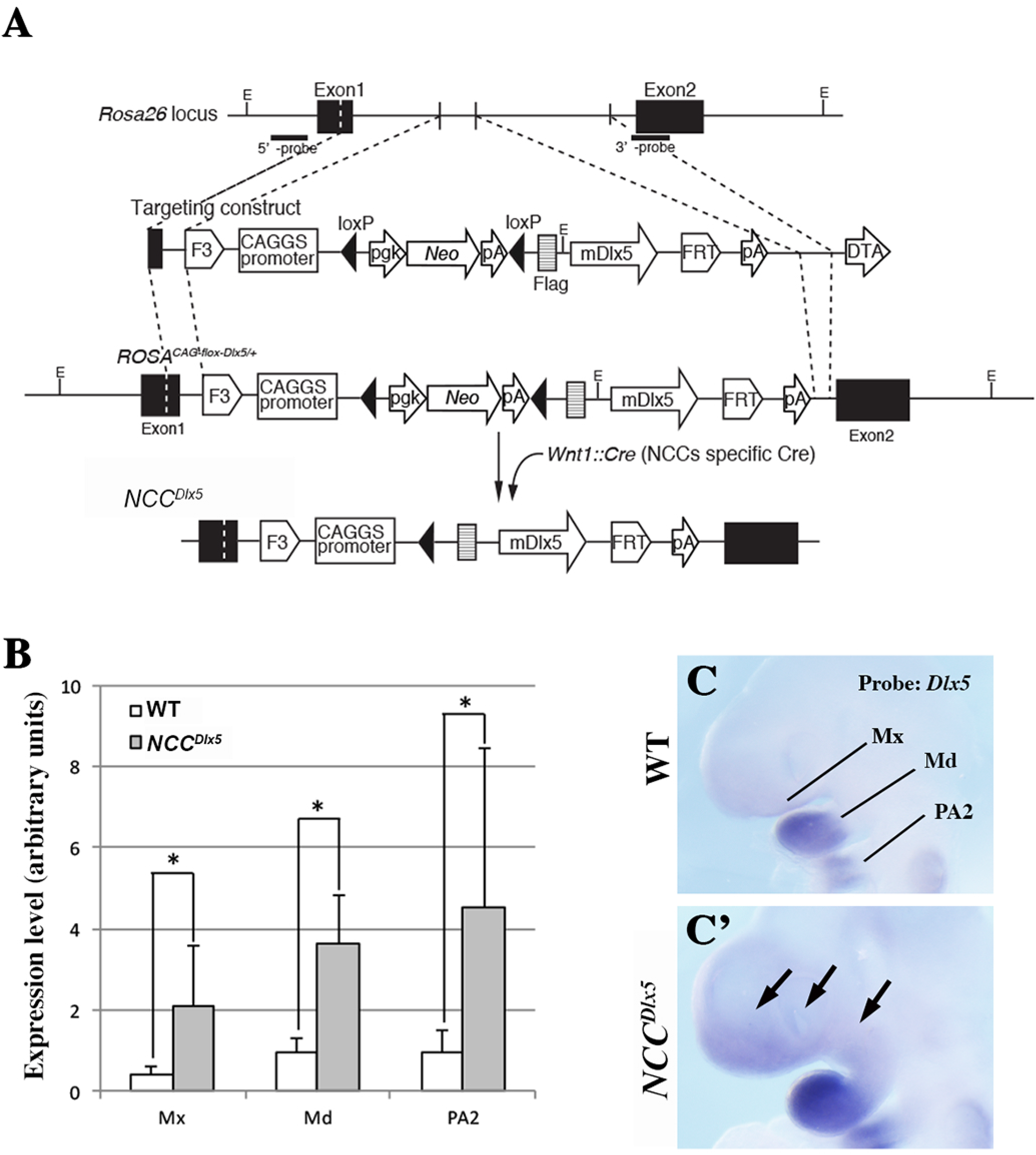
Generation of *NCC^Dlx5^* mice. A) Strategy for conditional expression of *Dlx5* from the *Rosa26* locus and generation of *NCC^Dlx5^* mice. Probes for genotyping are indicated as 5’- and 3’-probes. E, *Eco*RI. B) Comparison of *Dlx5* mRNA levels in the maxillary arch (Mx), mandibular arch (Md) and PA2 of control and *NCC^Dlx5^* E10.5 embryo. Messenger RNA levels were estimated by qRT-PCR. The values showed on the graph were mean ± SD of 5 duplicated samples. Statistics: Mann-Whitney U test using R-software (version 3.1.3). *p<0.01. C, C’) Whole mount *in situ* hybridization for *Dlx5* on control and *NCC^Dlx5^* E9.5 mice. Black arrows: areas of ectopic *Dlx5* expression.

To inactivate *Dlx5/6* in NCCs, *Dlx5/6^flox/flox^* mice, in which the homeodomain-encoding regions of both *Dlx5* and *Dlx6* are flanked by *lox* sequences (Bellessort, et al., 2016), were crossed with *Wnt1^Cre-ERT2/+^* mice in which tamoxifen exposure induces Cre-recombinase activity in cells of the developing neural tube, in migrating NCCs and in dopaminergic neurons, but not in other cell types. To cover most of the period of neural crest delamination and migration (Danielian, et al., 1998) *Dlx5/6^flox-flox^*::*Wnt1^Cre-ERT2/+^* timed pregnant dams received two intraperitoneal injections of tamoxifen at embryonic developmental days E6 and E7, generating *NCC*^Δ*Dlx5/6*^ embryos. The need to inactivate both *Dlx5* and *Dlx6* in NCCs derives from the fact that these two closely related genes are redundant in defining lower jaw identity: both need to be inactivated to transform the identity of the lower jaw into an upper jaw (Beverdam, et al., 2002)’ (Depew, et al., 2002).

### Craniofacial defects observed after deregulation of *Dlx5/6* expression in NCCs

At E18.5, *NCC^Dlx5^* foetuses presented a fully penetrant phenotype characterized by a short snout, open eyelids, misaligned vibrissae (Fig. 2A-B”) correctly located in the distal territory of the snout, and a cleft palate with no obvious signs of palatine rugae (Fig. 2A”’, B”’; Fig. 3D”; Fig. 5B”, C”). On the contrary, E18.5 *NCC*^Δ*DL·5/6*^ foetuses (Fig. 2C, C’) presented a marked mandibular retrognatia, with vibrissae present on both upper and lower jaws. In the upper jaw of *NCC*^Δ*Dlx5/6*^ foetuses, vibrissae were arranged in 5 rows as in control littermates, while ectopic vibrissae in the lower jaw were very close to each other and did not present a recognizable pattern; in these animals some vibrissae developed close to the midline. The palate and the eyelids of E18.5 *NCC*^Δ*Dlx5/6*^ foetuses did not present any obvious malformation.

**Figure 2:**
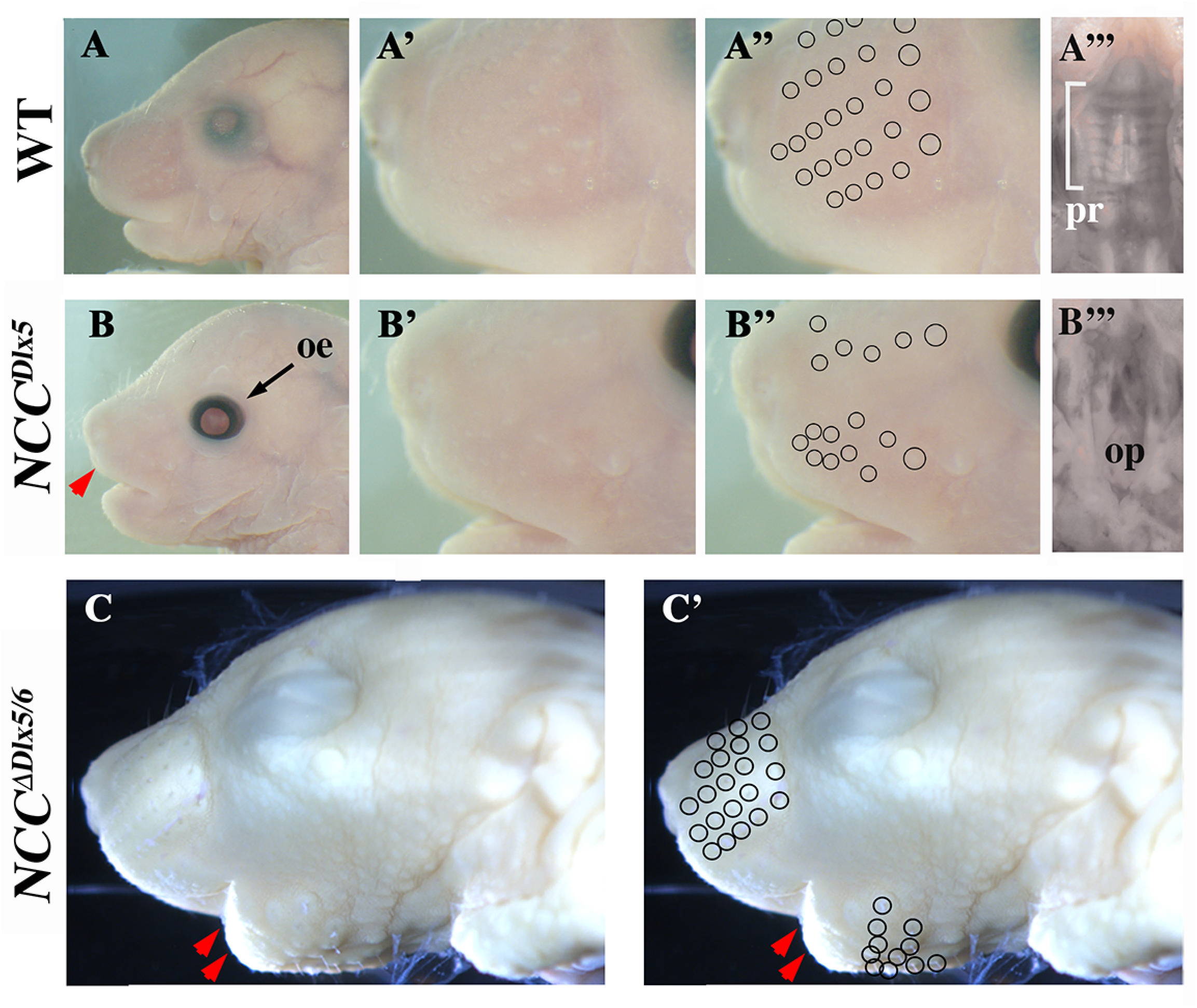
External appearances of *NCC^Dlx5^* and *NCC*^Δ*Dlx5/6*^ mouse heads at E18.5. Craniofacial appearance of control (A-A”’), *NCC^Dlx5^* (B-B”’) and *NCC*^Δ*Dlx5/6*^ (C, C’) mice. *NCC^dìx5^* mice have open eyelids (oe), a shortened snout (red arrowhead in B), misaligned vibrissae (open circles in A”, B”) and an open palate (op in B”’) with no sign of palatine rugae (pr in A”’). *NCC*^Δ*Dlx5/6*^ mouse heads present a severe mandibular retrognatia (red arrows in C, C’) with vibrissae well-aligned in the upper jaw, but also appearing ectopically in the lower jaw (open circles in C’).

The craniofacial phenotypes resulting from either up- or down-regulation of *Dlx5/6* in NCCs were analysed by 3D reconstruction after serial sectioning and imaging of the heads at E18.5 and compared to normal littermates or to heads of foetuses of the same age in which *Dlx5* and *Dlx6* had been constitutively inactivated (*Dlx5/6^-/-^* foetuses) (Gitton, et al., 2014; Beverdam, et al., 2002) (Figs. 3, 4 and Sup. Figs. 1-4 for 3D pdf files). As already described (Gitton, et al., 2014; Beverdam, et al., 2002; Depew, et al., 2002), after constitutive *Dlx5/6* inactivation, both lower and upper jaws are transformed resulting in a symmetric mouth. In *Dlx5/6^-/-^* mice, the lower jaw acquires morphological and molecular hallmarks of an upper jaw (Beverdam, et al., 2002)’(Depew, et al., 2002)’(Jeong, et al., 2008) while the premaxillary bone is not recognizable (Fig. 3 B-B”). Inactivation of *Dlx5/6* only in NCCs (*NCC*^Δ*Dlx5/6*^ embryos) resulted in severely reduced and virtually unrecognizable dentary bone, while maxillary bones, palate and premaxillary bone presented a relatively normal morphology (Fig. 3C’, C”). At variance with what observed in control and *Dlx5/6^-/-^* embryos, the transformed dentary bones of *NCC*^Δ*Dlx5/6*^ mice were fused along the midline and presented medio-lateral processes with a general structure reminiscent of maxillary bones and palate. The *NCC*^Δ*Dlx5/6*^ transformed dentary supported two lower incisors which, in *Dlx5/6^-/-^* embryos, are not enclosed in the transformed bone. Although morphologically distinct form the upper jaws, the lower jaws of *NCC*^Δ*Dlx5/6*^ embryos presented further hallmarks of maxillary identity with well developed vibrissae (Fig. 2C, C’; Fig. 5B, B’) and absence of a recognizable Meckelian cartilage (Fig. 4C, C’; Fig. 5C, C’). Remarkably the infraorbital foramen, a major trait of the mammalian maxilla was present in both the upper and lower jaws of *Dlx5/6^-/-^* and *NCC*^Δ*Dlx5/6*^ embryos (Sup. Figs. 1-3) reinforcing the notion of a switch in identity.

**Figure 3:**
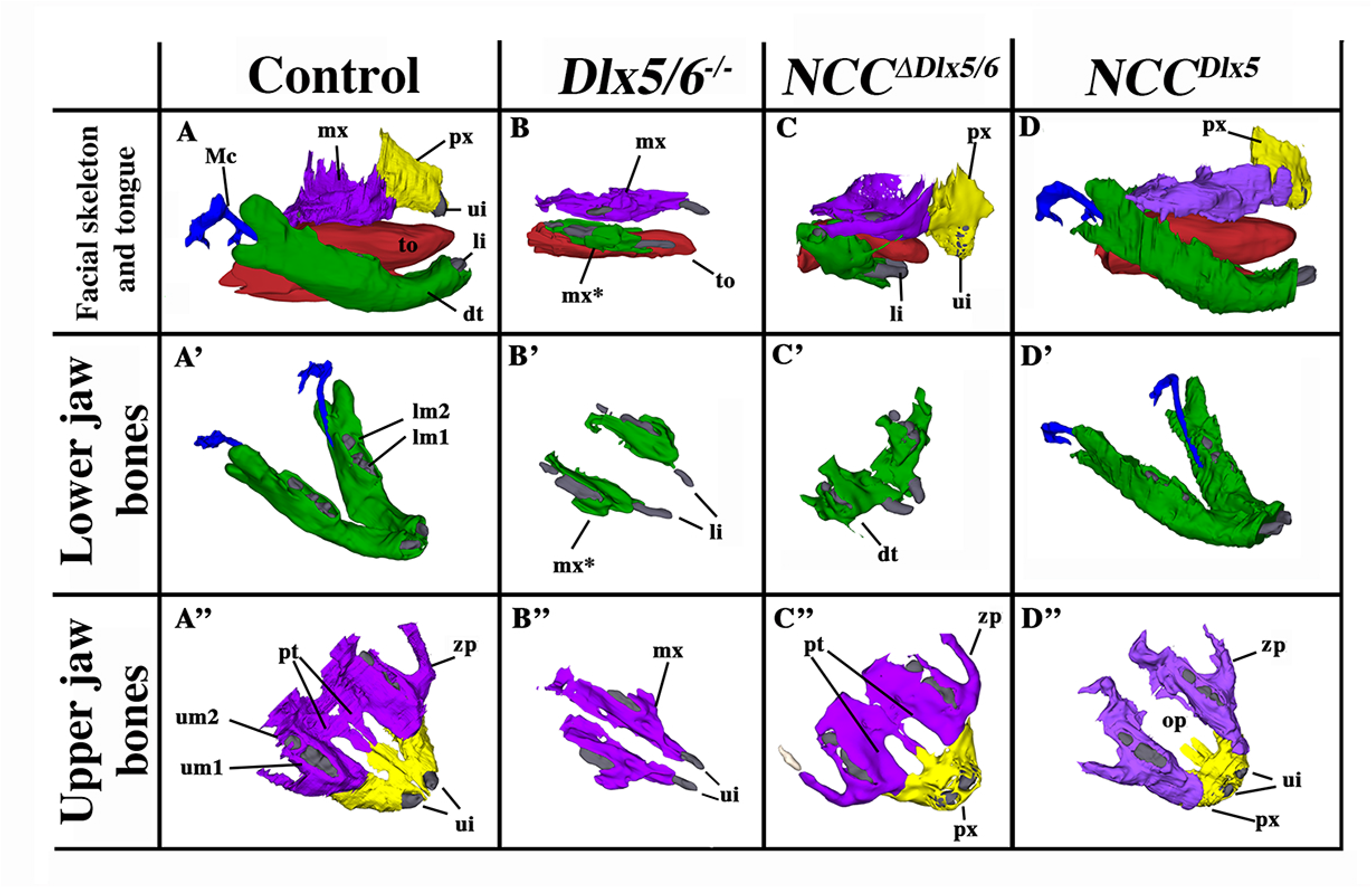
Selected views of bone and cartilagineous elements from 3D reconstructions of control, *Dlx5/6^-/-^*, *NCC^mix5/6^* and *NCC^Dlx5^* E18.5 mouse foetuses. A-A”) Skeletal elements from normal mouse embryonic heads. B-B”) After constitutive *Dlx5/6* inactivation both upper and the lower jaw are transformed generating similar bony elements and a highly symmetric mouth. Incisors (li, ui) are short and not in direct contact with the transformed dentary (mx*) or maxillary (mx) bones. The premaxillary bone is absent. C-C”) *NCC*^Δ*Dlx5/6*^ heads present a severely shortened dentary bone (green in C, C’) which fuses in the midline and displays well developed lateral processes reminiscent of maxillary bones. Maxillary and premaxillary (px) bones are relatively well-developed with a closed palate (pt) and normal zygomatic processes (zp). D-D”) *NCC^Dlx5^* mice present an almost normal dentary bone with a reduced coronoid process (see Fig. 6) and well developed Meckelian cartilage (D’). In *NCC^Dlx5^* mice maxillary bones are abnormal, they present an open palate (op), and protrude anteriorly, the zygomatic process of the maxilla (zp) is malformed, the premaxillary bone (px) is reduced. Abbreviations: dt dentary bone; li, lower incisor; lm1, lm2, lower molars 1 and 2; Mc, Mekelian cartilage; mx, maxillary bones; mx*, transformed dentary bone; op, open palate; pt, palate; px, premaxillary bone; ui, upper incisor; to, tongue; zp, zygomatic process.

**Figure 4:**
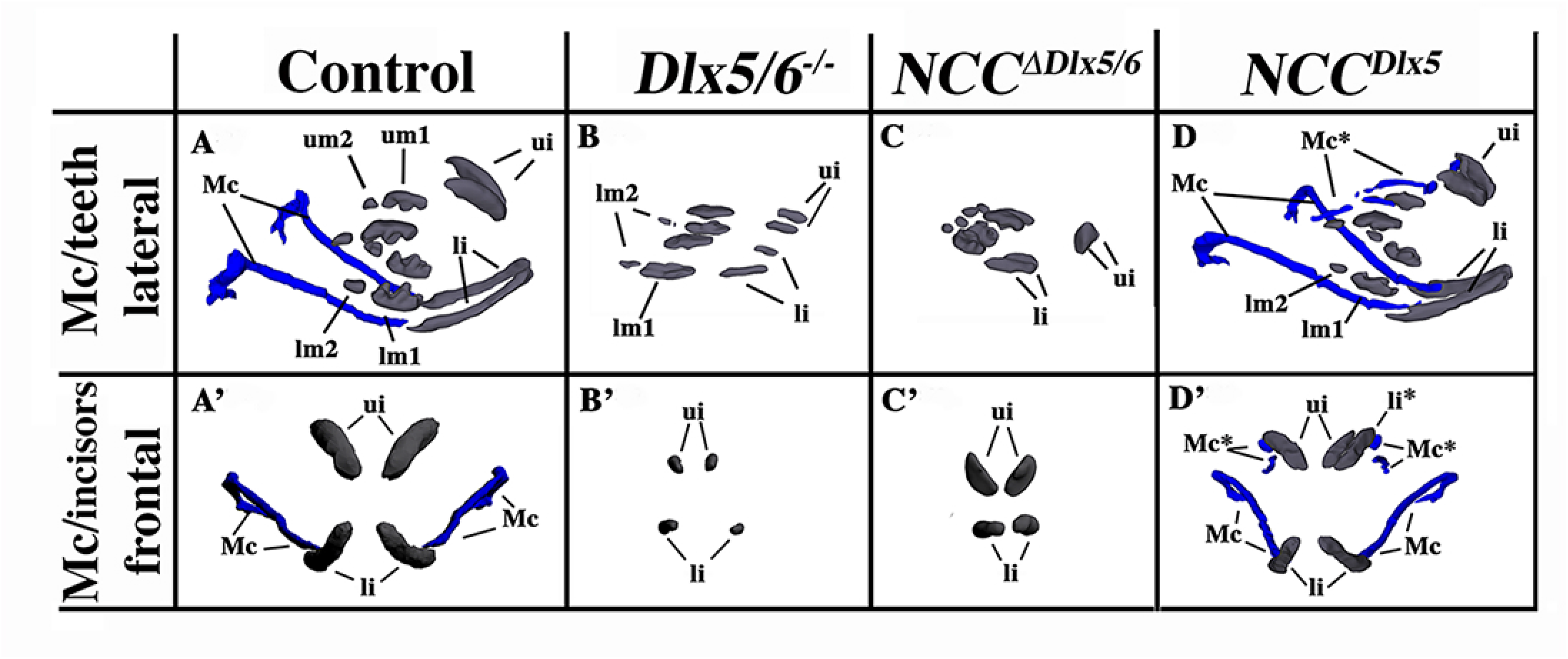
Teeth complement and Meckelian cartilage from 3D reconstructions of control, *Dlx5/6^-/-^*, *NCC^∆Dlx5/6^* and *NCC^Dlx5^* E18.5 mouse foetuses. Lateral (A-D) and frontal (A’-D’) views of teeth (grey) and Mekelian cartilage (blue) from control (A, A’), *Dlx5/6^f^-* (B, B’), *NCC^ΔDL·5/6^* (C, C’) and *NCC^Dlx5^*E18.5 mouse foetuses. The Meckelian cartilage (Mc) is absent in *Dlx5/6^-/-^* and *NCC*^Δ*Dlx5/6*^ foetuses and is well formed in *NCC^Dlx5^* foetuses where supernumerary Meckel-like cartilage bars are present in the upper jaw (Mc*). In *Dlx5^-/-^* foetuses the incisors (ui, li) are short and straight and are not supported by bony elements. In *NCC*^Δ*Dlx5/6*^ foetuses the incisors are also short, but converge towards the midline while in *NCC^Dlx5^* foetuses lower incisor are apparently normal while upper incisors are longer than normal and often duplicated with the second incisor (li*) supported by the transformed maxillary bone suggesting, therefore, that it might represent a transformed lower incisor. Abbreviations: li, lower incisor; lm1, lm2, lower molars 1 and 2; li*, transformed upper incisor; Mc, Meckelian cartilage; Mc*, duplicated Meckelian cartilage; ui, upper incisor.

**Figure 5:**
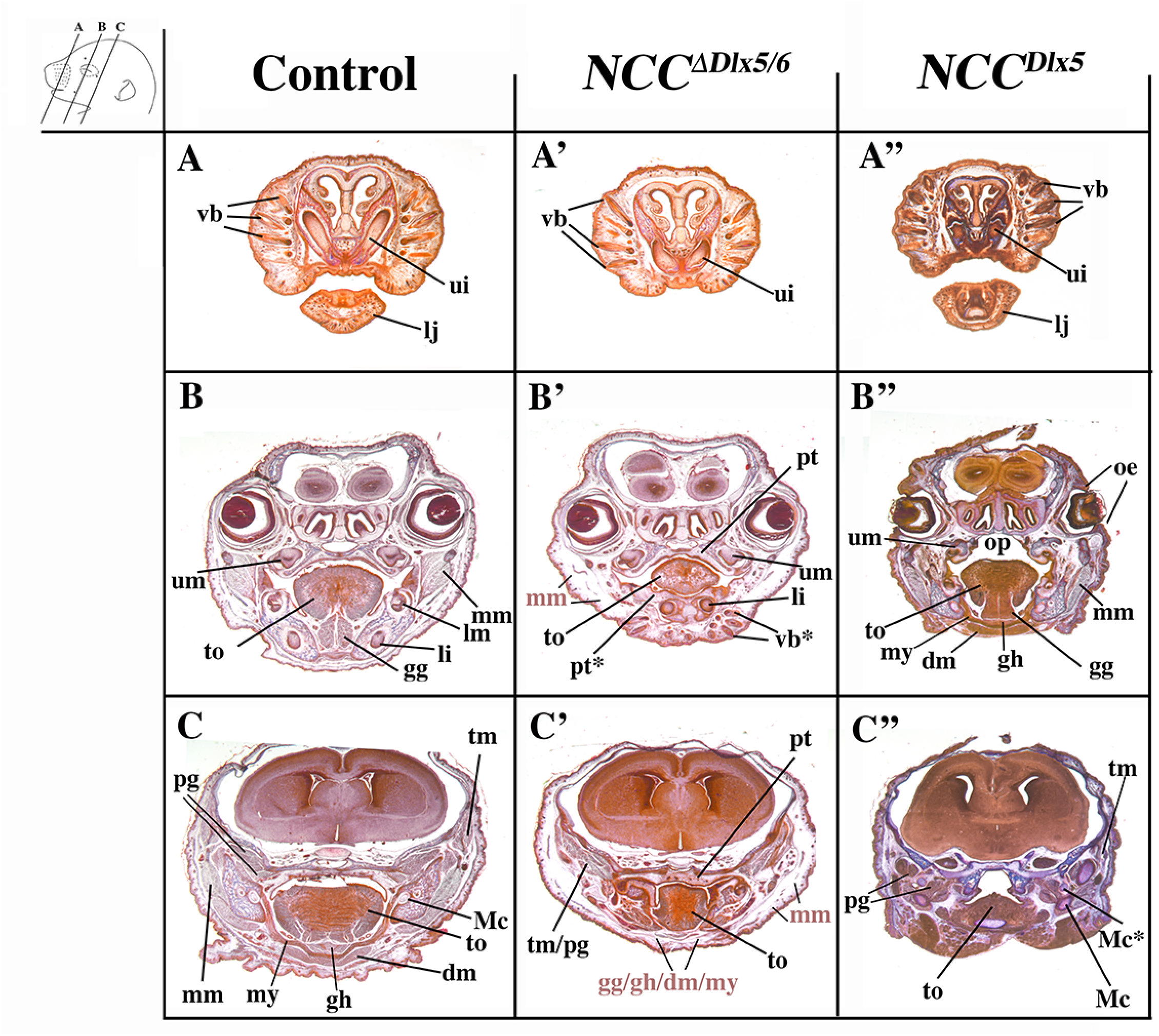
Representative cranial frontal sections of E18.5 control, *NCC^∆Dlx5/6^* and *NCC^Dlx5^* mouse foetuses. Compared to control foetuses (A-C), *NCC*^Δ*Dlx5/6*^ foetuses (A’-B’) present ectopic vibrissae (vb*) in the lower jaw (B’), reduced and disorganized tongue musculature (to), absence of masseter, digastric, mylohyoid, geniohyoid and genioglossus muscles while the temporal and pterygoid muscles (tm, pg) form but show defective attachments on the transformed lower jaws. The midline fusion of the transformed dentary bone of *NCC*^Δ*Dlx5/6*^ foetuses gives rise to a palate-like structure in the lower jaw (pt*) with folds reminiscent of palatine rugae. *NCC^Dlx5^*foetuses (A”-C”) present open eyelids (oe), an open palate (op), ectopic Meckelian-like cartilages, relatively normal tongue and associated musculature and mispatterned masticatory muscles adapted to the transformed skeletal elements. Planes of section are indicated in the upper left insert. Abbreviations: dm, digastric muscle; gg, genioglossus muscle; gh, geniohyoid muscle; li, lower incisor; lm, lower molar; lj, lower jaw; Mc, Mekelian cartilage; Mc*, duplicated Mekelian cartilage; mm, masseter muscle; mx maxillary bones; mx*, transformed dentary bone; my, mylohyoid muscle; oe, open eyelids; op, open palate; pg, lateral and medial pterygoid muscles; pt palate; pt*, ectopic palate-like structure resulting from midline fusion of lower jaws; px, premaxillary bone; ui, upper incisor; um, upper molar; tm, temporal muscle; to, tongue; vb, vibrissae; vb*, ectopic vibrissae. Abbreviations in light red represent muscles that did not differentiate and are replaced by loose mesenchymal tissue.

*NCC^Dlx5^* mice presented a relatively normal lower jaw, with a well-developed Meckelian cartilage and dentary bone (Figs. 3D, D’; 4D, D’; 6A-B’). Maxillary bones were hypomorphic with an open palate and smaller zygomatic and premaxillary bones (Figs. 3D”; Fig. 5B”, C”; Fig. 6). We also observed the presence of fragmented cartilaginous rods converging towards the midline of the upper jaw (Fig. 4D, D’; Fig. 5C”) suggesting the presence of an ectopic Meckelian cartilage. Skeletal preparations of E18.5 *NCC^Dlx5^* foetuses (Figs. 6; 7) confirmed the malformation of upper jaw bones and cartilage components and the relatively normal appearance of lower jaw elements except for shortening of the coronoid process of the dentary bone (Fig. 6B, B’). The premaxillary bone was reduced (Fig. 3A”, D”; Fig. 6A, A’), the zygomatic process of the maxilla was thicker and shorter (Fig. 6C, C’), the jugal bone was shorter (Fig. 6A, A’; C, C’) and the zygomatic and retroarticular processes of the squamosal were missing (Fig. 6C, C’). In *NCC^Dlx5^* embryos the incus seemed to have partially acquired a malleus-like identity as: 1) it short process was elongated, 2) a small ectopic bone, which could be interpreted as a duplicated gonial bone, appeared adjacent to the incus, and 3) the malleus-incus joint, which is normally of ball-and-socket type, was symmetric (Fig. 6D, D’; Sup. Fig. 5). The infraorbital foramen was not present in *NCC^Dlx5^* mice also suggesting an upper jaw transformation (Sup. Fig. 4).

**Figure 6:**
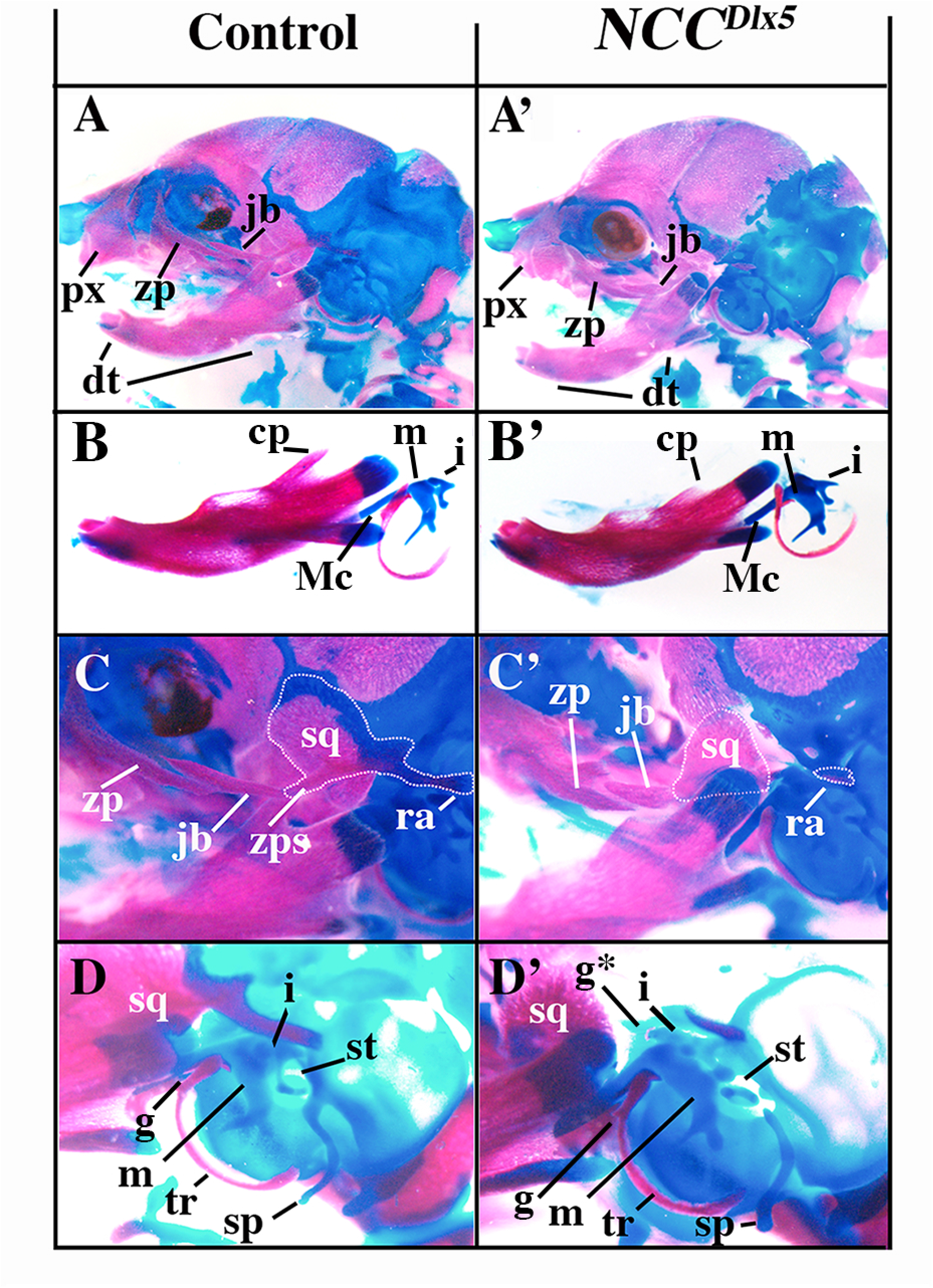
Craniofacial skeletal preparations from control and *NCC^Dlx5^* E18.5 foetuses. (A, A’) Lateral views of the head of control (A) and *NCC^Dlx5^* (A’) E18.5 foetuses. In *NCC^Dlx5^*mice the maxilla is deformed with thickening of the zygomatic process while the jugal and the premaxillary bones are shortened. (B, B’) Dentary bones of control (B) and *NCC^Dlx5^* (B’) E18.5 foetuses. The coronoid process (cp) is hypoplastic in *NCC^Dlx5^* mice while the Meckelian cartilage (Mc) and the malleus (m) are well-formed. (C, C’) Jaw joint region of control (C) and *NCC^Dlx5^* (C’) E18.5 mouse foetuses. The zygomatic and retroarticular processes (ra) of the squamosal (sq) are missing and the coronoid process of the dentary bone is hypoplastic in *NCC^Dlx5^* mice. (D, D’) Middle ear components of control (D) and *NCC^Dlx5^* (D’) E18.5 mouse foetuses. The incus (i) is deformed and dislocated from the stapes (st) fused to the styloid process (sp) in *NCC^Dlx5^* mice. Abbreviations: cp, coronoid process; dt, dentary bone; i, incus; jb, jugal bone; m, malleus; Mc, Meckel’s cartilage; pmx, premaxilla; ra, retroarticular process of squamosal; sp, styloid process; sq, squamosal; st, stapes; tr, ectotympanic ring; zp, zygomatic process of maxilla; zps, zygomatic process of squamosal.

**Figure 7:**
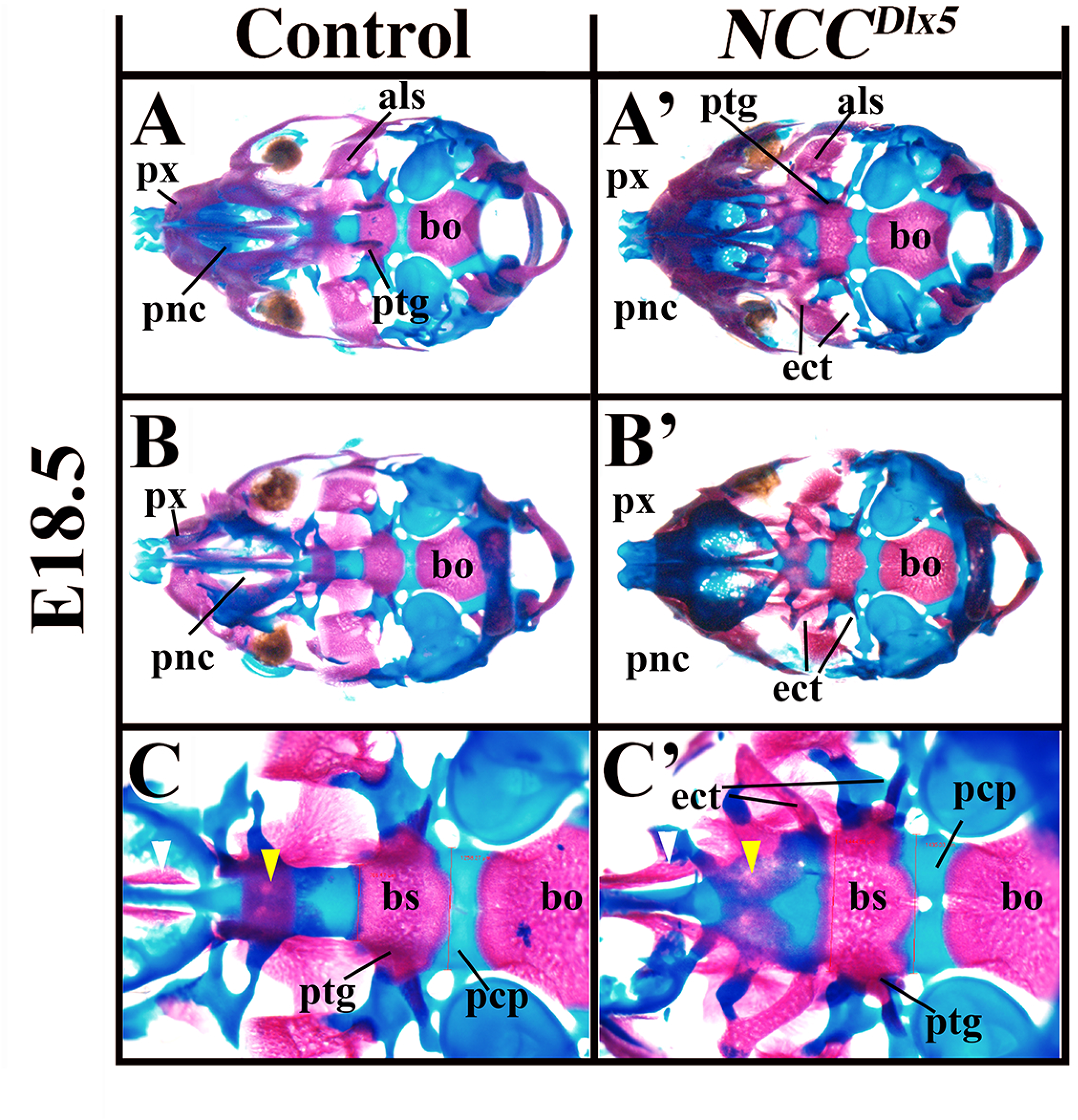
Cranial base deformity in NCCDlx5 mice. Ventral (A, A’; C, C’) and dorsal (B, B’) views of the cranial base of E18.5 control (A, B, C) and *NCC^Dlx5^* (A’, B’, C’) E18.5 mouse foetuses. In *NCC^Dlx5^* mice, palatal processes of the maxilla (white arrowhead in C, C’) and palatine (yellow arrowhead in C, C’) are defective. The paranasal cartilage (pnc), basisphenoid (bs) and the prechordal plate (pcp) are enlarged in width. Ectopic cartilaginous and osseous struts extended from the basisphenoid laterally and anteriorly, respectively (ect). Abbreviations: als, alisphenoidal bone; bo, basioccipital bone; bs, basisphenoidal bone; ect, ectopic structure; pcp, parachordal plate; px, premaxilla; pnc, paranasal cartilage; ptg, pterygoid.

### Incisor phenotypes

Lower incisor germs of normal E18.5 foetuses are elongated structures that develop within the dentary bone and converge medially towards the distal tip (Fig. 4A, A’ and Sup. Fig. 1). Normal lower incisors form an occlusive pattern with upper incisor germs, which are supported by the premaxillary bone. In normal animals, upper incisors are much shorter and thicker than their lower jaw counterpart and also converge towards the midline (Fig. 4A, A’ and supplementary Figure 1). Remarkably, in E18.5 *Dlx5/6^-/-^* foetuses, the four incisor germs were not supported by any bone structure (dentary or premaxillary) and appeared as short rods of comparable length oriented along the proximo-distal axis not converging toward the midline (Figs. 3B-B”; 4B, B’ and Sup. Fig. 2). The four incisors germs of *Dlx5/6^-/-^* foetuses were, therefore, very similar to each other. The lack of supporting bones for *Dlx5/6^-/-^* incisors suggests that bones and incisors teeth can develop independently. In *NCC*^Δ*Dlx5/6*^ E18.5 foetuses (Figs. 3C’, 4C, C’ and Sup. Fig. 3), lower incisors were constituted by short, straight rods oriented towards the midline and protruding for about two thirds of their length from the transformed dentary bone. Upper incisors germs of *NCC*^Δ*Dlx5/6*^ mutants were short and pointed structures entirely supported by premaxillary bones; they were oriented ventrally along an oblique axis towards the midline.

Incisor morphology of *NCC^Dlx5^* E18.5 foetuses was particularly interesting (Fig. 4D, D’ and Supplementary Fig. 4). Lower incisors were apparently normal in shape and position; upper incisors were often duplicated: a relatively normal short upper incisor was supported by the premaxillary bone and was juxtaposed to a second, longer, upper incisor that was partially supported by the transformed maxillary bone (to visualize the site of insertion see Supplementary Fig. 4). When a single upper incisor was found, it was larger on the medio/lateral axis suggesting that it might represent the fusion of two parallel incisors. This finding could further support the view that over expression of *Dlx5* in maxillary NCCs results in the transformation of maxillary into mandibular structures carrying an incisor.

### Ectopic *Dlx5* expression in NCCs affects the development of cranial base structures

*NCC^Dlx5^* mice presented changes in the cranial base, which is partially derived from NCCs (McBratney-Owen, et al., 2008; Jiang, et al., 2002); on the contrary, *NCC*^Δ*Dlx5/6*^ foetuses did not present obvious differences of these structures. In *NCC^Dlx5^* mice, midline structures of the cranial base including the paranasal cartilage, the presphenoid and the basisphenoid were larger than in control mice (Fig. 7A-C’). From the basisphenoid, ectopic cartilaginous and osseous struts extended laterally and anteriorly; the later ectopic process fused to the hypochiasmatic cartilage of mesodermal origin, disturbing the formation of the optic foramen (Fig. 7C, C’).

### *Dlx5/6* expression in neural crest cells is required for proper jaw muscularization

We have previously shown that *Dlx5/6^-/-^* mice display agenesis of masticatory muscles due to the lack of instructive cues from NCCs onto cephalic myogenic mesoderm precursors (Heude, et al., 2010). In E18.5 *NCC*^Δ*Dlx5/6*^ foetuses, craniofacial muscularization was affected with a phenotype strongly resembling that observed in *Dlx5/6^-/-^* mice (Heude, et al., 2010). The masticatory masseter muscles failed to differentiate normally and were replaced by loose mesenchymal tissue (Fig.5B’). The other masticatory muscles, temporal and pterygoids, were present dorsally but showed defective attachments on the transformed lower jaws (Fig. 5C’). The tongue and associated musculature were severely affected: the suprahyoid muscles (mylohyoid, digastric and geniohyoid) and the extrinsic genioglossus tongue muscles did not differentiate and were replaced by loose tissue containing few disorganized myofibers; the intrinsic tongue musculature was reduced, disorganized and remained as a vestigial medially-located structure disconnected from lower jaw bones (Fig. 5B’, C’).

In *NCC^Dlx5^* foetuses all craniofacial muscles could be identified, but their attachment points and their shape were changed to adapt to the transformed skeletal elements (Fig. 5 B”, C”). These observations reinforce the notion that cephalic myogenic precursors require *Dlx5/6*-dependent morphogenetic instructions from NCCs for proper jaw muscle patterning and differentiation (Sugii, et al., 2017).

### *Dlx5* overexpression in NCCs switches maxillary to mandibular transcriptomic signature

To identify regulatory pathways involved in determining jaw identity, we performed transcriptome analysis on PA tissues of E10.5 control and *NCC^Dlx5^* embryos. When the *NCC^Dlx5^* maxillary arch samples were compared to those from control maxillary arches, 12 and 21 genes were identified as increased or decreased by more than 2-fold, respectively (Table 1). Scatter plot of the signal intensity showed that the upregulated genes corresponded to mandibular marker previously reported to be downstream of Dlx5/6 (Fig. 8) (Jeong, et al., 2008). By contrast, maxillary marker genes reported to be upregulated in the transformed mandibular arch of *Dlx5/6^-/-^* embryos showed only small deviation from the diagonal (Fig. 8). To confirm this difference and further characterize the Dlx5-downstream genes, we then performed microarray analysis on PA tissues of E10.5 *Dlx5/6^-/-^* embryos and compared the results with genes deregulated in the *NCC^Dlx5^* maxillary arch. In the *Dlx5/6^-/-^* mandibular arch we identified 18 genes whose expression was increased and 45 genes whose expression was decreased by more than 2-folds (Table 2). Remarkably, 10 of 12 genes up-regulated in the *NCC^Dlx5^* maxillary arch were also down-regulated in the *Dlx5/6^-/-^* mandibular arch suggesting a common set of downstream targets involved in lower jaw specification (Table 2). By contrast, only 1 gene, *Has2*, a putative hyaluronan synthase, was shared between the up-regulated genes in *Dlx5/6^-/-^* mandibular arch and the down-regulated genes in *NCC^Dlx5^*maxillary arch (Table 2).

**Figure 8:**
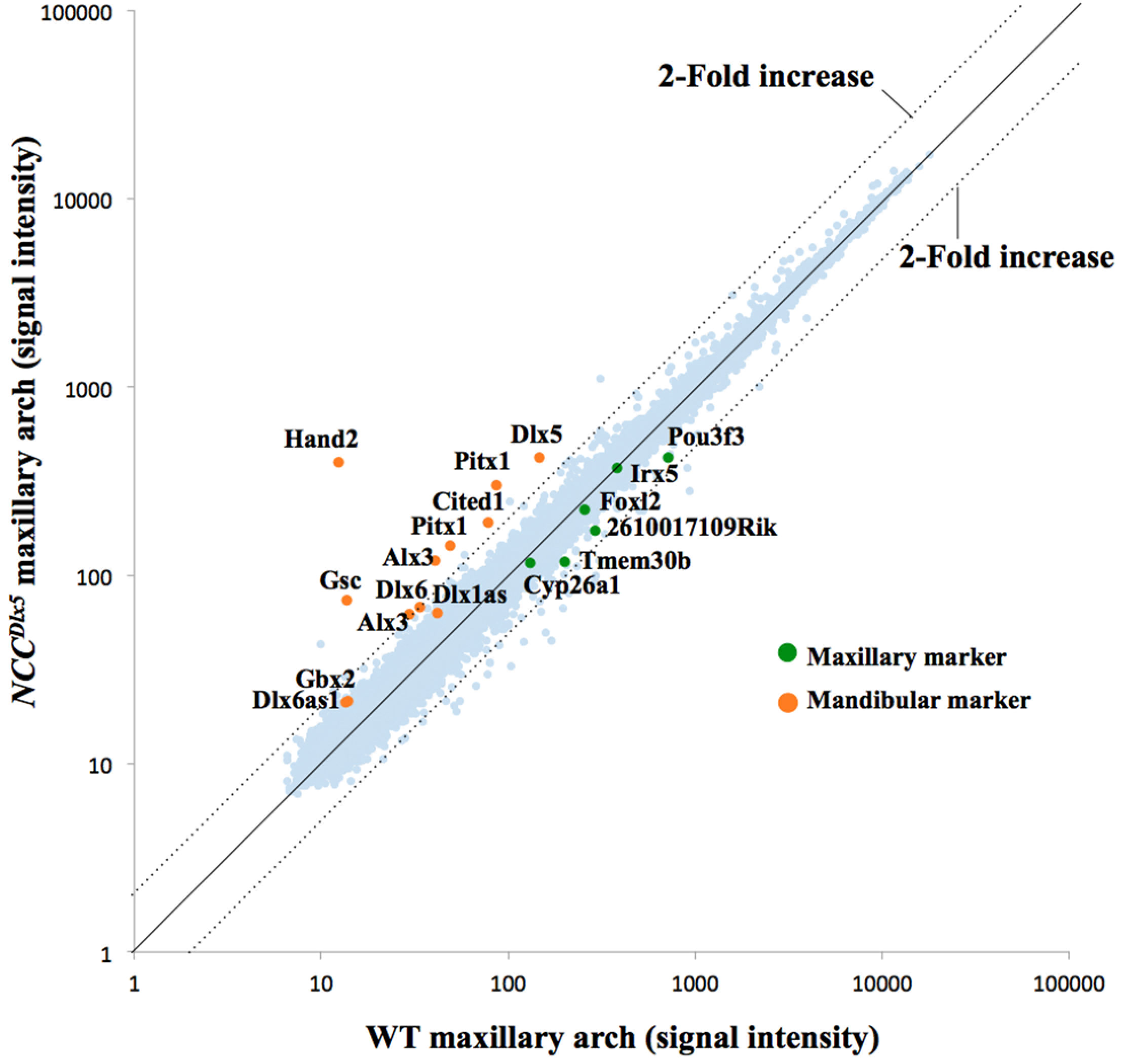
Scatter plot of signal intensities representing differential gene expression in the E10.5 control and *NCC^Dlx5^* maxillary arches. Points corresponding to known maxillary and mandibular arch markers are colored green and orange, respectively.

**TABLE 1:**
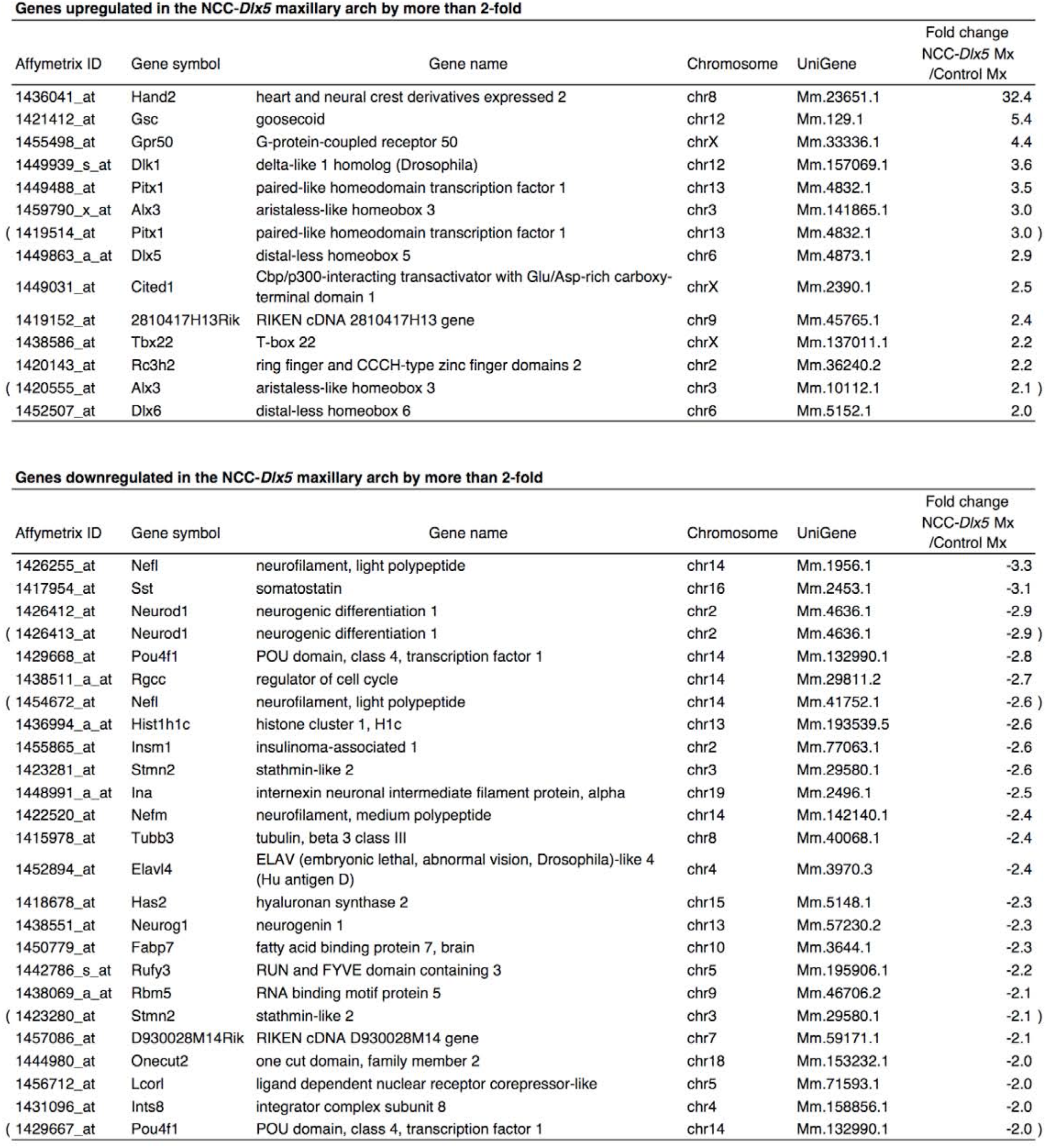
Genes up- or down-regulated in the *NCCDlx5* maxillary arch.

| Affymetrix ID | Gene symbol | Gene name | Chromosome | UniGene | Fold change<br>NCC-*Dlx5* Mx<br>/Control Mx |
| --- | --- | --- | --- | --- | --- |
| 1436041_at | Hand2 | heart and neural crest derivatives expressed 2 | chr8 | Mm.23651.1 | 32.4 |
| 1421412_at | Gsc | goosecoid | chr12 | Mm.129.1 | 5.4 |
| 1455498_at | Gpr50 | G-protein-coupled receptor 50 | chrX | Mm.33336.1 | 4.4 |
| 1449939_s_at | Dlk1 | delta-like 1 homolog (Drosophila) | chr12 | Mm.157069.1 | 3.6 |
| 1449488_at | Pitx1 | paired-like homeodomain transcription factor 1 | chr13 | Mm.4832.1 | 3.5 |
| 1459790_x_at | Alx3 | aristaless-like homeobox 3 | chr3 | Mm.141865.1 | 3.0 |
| ( 1419514_at | Pitx1 | paired-like homeodomain transcription factor 1 | chr13 | Mm.4832.1 | 3.0 ) |
| 1449863_a_at | Dlx5 | distal-less homeobox 5 | chr6 | Mm.4873.1 | 2.9 |
| 1449031_at | Cited1 | Cbp/p300-interacting transactivator with Glu/Asp-rich carboxy-terminal domain 1 | chrX | Mm.2390.1 | 2.5 |
| 1419152_at | 2810417H13Rik | RIKEN cDNA 2810417H13 gene | chr9 | Mm.45765.1 | 2.4 |
| 1438586_at | Tbx22 | T-box 22 | chrX | Mm.137011.1 | 2.2 |
| 1420143_at | Rc3h2 | ring finger and CCCH-type zinc finger domains 2 | chr2 | Mm.36240.2 | 2.2 |
| ( 1420555_at | Alx3 | aristaless-like homeobox 3 | chr3 | Mm.10112.1 | 2.1 ) |
| 1452507_at | Dlx6 | distal-less homeobox 6 | chr6 | Mm.5152.1 | 2.0 |

Genes downregulated in the *NCC-Dlx5* maxillary arch by more than 2-fold
| Affymetrix ID | Gene symbol | Gene name | Chromosome | UniGene | Fold change<br>NCC-*Dlx5* Mx<br>/Control Mx |
| --- | --- | --- | --- | --- | --- |
| 1426255_at | Nefl | neurofilament, light polypeptide | chr14 | Mm.1956.1 | -3.3 |
| 1417954_at | Sst | somatostatin | chr16 | Mm.2453.1 | -3.1 |
| 1426412_at | Neurod1 | neurogenic differentiation 1 | chr2 | Mm.4636.1 | -2.9 |
| ( 1426413_at | Neurod1 | neurogenic differentiation 1 | chr2 | Mm.4636.1 | -2.9 ) |
| 1429668_at | Pou4f1 | POU domain, class 4, transcription factor 1 | chr14 | Mm.132990.1 | -2.8 |
| 1438511_a_at | Rgcc | regulator of cell cycle | chr14 | Mm.29811.2 | -2.7 |
| ( 1454672_at | Nefl | neurofilament, light polypeptide | chr14 | Mm.41752.1 | -2.6 ) |
| 1436994_a_at | Hist1h1c | histone cluster 1, H1c | chr13 | Mm.193539.5 | -2.6 |
| 1455865_at | Insm1 | insulinoma-associated 1 | chr2 | Mm.77063.1 | -2.6 |
| 1423281_at | Stmn2 | stathmin-like 2 | chr3 | Mm.29580.1 | -2.6 |
| 1448991_a_at | Ina | internexin neuronal intermediate filament protein, alpha | chr19 | Mm.2496.1 | -2.5 |
| 1422520_at | Nefm | neurofilament, medium polypeptide | chr14 | Mm.142140.1 | -2.4 |
| 1415978_at | Tubb3 | tubulin, beta 3 class III | chr8 | Mm.40068.1 | -2.4 |
| 1452894_at | Elavl4 | ELAV (embryonic lethal, abnormal vision, Drosophila)-like 4 (Hu antigen D) | chr4 | Mm.3970.3 | -2.4 |
| 1418678_at | Has2 | hyaluronan synthase 2 | chr15 | Mm.5148.1 | -2.3 |
| 1438551_at | Neurog1 | neurogenin 1 | chr13 | Mm.57230.2 | -2.3 |
| 1450779_at | Fabp7 | fatty acid binding protein 7, brain | chr10 | Mm.3644.1 | -2.3 |
| 1442786_s_at | Rufy3 | RUN and FYVE domain containing 3 | chr5 | Mm.195906.1 | -2.2 |
| 1438069_a_at | Rbm5 | RNA binding motif protein 5 | chr9 | Mm.46706.2 | -2.1 |
| ( 1423280_at | Stmn2 | stathmin-like 2 | chr3 | Mm.29580.1 | -2.1 ) |
| 1457086_at | D930028M14Rik | RIKEN cDNA D930028M14 gene | chr7 | Mm.59171.1 | -2.1 |
| 1444980_at | Onecut2 | one cut domain, family member 2 | chr18 | Mm.153232.1 | -2.0 |
| 1456712_at | Lcorl | ligand dependent nuclear receptor corepressor-like | chr5 | Mm.71593.1 | -2.0 |
| 1431096_at | Ints8 | integrator complex subunit 8 | chr4 | Mm.158856.1 | -2.0 |
| ( 1429667_at | Pou4f1 | POU domain, class 4, transcription factor 1 | chr14 | Mm.132990.1 | -2.0 ) |

**TABLE 2:** Comparison of genes affected in the *NCC^Dlx5^* maxillary arch with those affected in the *Dlx5/6*-null mandibular arch.

| NCC-*Dlx5* Mx/Control Mx |  | *Dlx5/6*-null Md/Control Md |  |
| --- | --- | --- | --- |
| Up | Down | Up | Down |
| Hand2 | Nefl | Pou3f3 | Dlx5 |
| Gsc | Sst | Tmem30b | Dlx6 |
| Gpr50 | Neurod1 | 2900092D14Rik | Dlx6os1 |
| Dlk1 | Pou4f1 | Itih5 | Dlx1as |
| Pitx1 | Rgcc | B230214O09Rik | Gm2818 |
| Alx3 | Hist1h1c | Bdnf | Gsc |
| Dlx5 | Insm1 | Foxl2 | Hand2 |
| Cited1 | Stmn2 | Foxl2os | Gpr50 |
| 2810417H13Rik | Ina | 2610017I09Rik | Tbx22 |
| Tbx22 | Nefm | Igf1 | Gbx2 |
| Rc3h2 | Tubb3 | Cyp26a1 | Pitx1 |
| Dlx6 | Elavl4 | Crym | Dgkk |
|  | Has2 | Has2 | Col8a2 |
|  | Neurog1 | Six1 | Alx3 |
|  | Fabp7 | Mtap2 | Dkk2 |
|  | Rufy3 | Ptx3 | Sdpr |
|  | Rbm5 | Figf | Dlx6os2 |
|  | D930028M14Rik | Ebf2 | Rgs5 |
|  | Onecut2 |  | Dlx4 |
|  | Lcorl |  | Cited1 |
|  | Ints8 |  | Lrrc17 |
|  |  |  | Zadh2 |
|  |  |  | Shox2 |
|  |  |  | Aldh1a2 |
|  |  |  | Osr1 |
|  |  |  | Ptprz1 |
|  |  |  | Tshz1 |
|  |  |  | Dlk1 |
|  |  |  | Pcdh19 |
|  |  |  | Gm6958 |
|  |  |  | Nr5a2 |
|  |  |  | Tnnt1 |
|  |  |  | Nrk |
|  |  |  | A730090H04Rik |
|  |  |  | 9430047L24Rik |
|  |  |  | Pmp22 |
|  |  |  | A130040M12Rik |
|  |  |  | Rspo2 |
|  |  |  | Arg2 |
|  |  |  | Synpo2 |
|  |  |  | Pdgfrl |
|  |  |  | Bmper |
|  |  |  | 1110006E14Rik |
|  |  |  | Plcxd3 |
|  |  |  | Hist3h2ba |

Overlapping genes are colored correspondingly.

## DISCUSSION

In gnathostomes, feeding depends on muscularized, articulated jaws capable to support prehension, mastication and swallowing. In modern tetrapods, upper and lower jaws are, in general, distinct anatomical structures. Since many decades, however, it has been remarked that in basal reptiles and amphibians the upper and the lower jaws are essentially mirror images of each other (Romer, 1940). The lower jaw of contemporary species is constituted by two mobile dentary bones that form around an early rod-like Meckelian cartilage and converge towards the centre where they fuse at the mandibular symphysis. The stationary upper jaw is mostly constituted by dermal bones that fuse along the midline and extend medio-laterally to form the palate and the maxillary part of the face. During embryonic development both upper and lower jaws derive from the first pharyngeal arch (PA1) that is colonized by Hox-negative NCCs (Kontges and Lumsden, 1996; Trainor and Tam, 1995; Couly, et al., 1993) which give rise to most cartilages, bones and tendons of the jaws, while the associated musculature originates from the myogenic mesoderm. Endothelin-1 (Edn1) is a key signal at the origin of asymmetric jaw development. Binding of Edn1 to its receptor A (Ednra) in NCCs activates the expression of lower-jaw specific genes including *Dlx5/6* and *Hand2* (Sato, et al., 2008b). Remarkably targeted inactivation of either *Edn1*, *Ednra*, *Dlx5/6*or *Hand2* result in the transformation of the lower jaw into new structures that present key morphological hallmarks of an upper jaw such as, for example, the presence of vibrissae and the absence of Meckelian cartilage (Ruest, et al., 2004; Ruest, et al., 2003; Clouthier, et al., 1998). It must be pointed out, however, that each of these mutations generates different lower jaw morphologies. Targeted constitutive inactivation of *Dlx5/6* results in the simultaneous transformation of the upper and of the lower jaws giving rise to a symmetric mouth in which each of the four jaws (upper/lower, right/left) presents a morphology very similar to each of the others transposed along reflection planes (Beverdam, et al., 2002; Depew, et al., 2002). We have shown that the Dlx5/6-dependent patterning of the upper jaw derives from their transitory expression in an ectodermal, distal, signalling centre located in the lamboid-junction area and not from their expression in mandibular NCCs (Gitton, et al., 2014). Constitutive disruption of either 1) *Edn1*, 2) its receptor *Ednra* in NCCs or 3) the Edn1 downstream target *Hand2* results in jaw phenotypes very different from those observed in *Dlx5/6^-/-^* embryos: the upper jaw does not present any obvious malformation while the lower jaw becomes small and hypomorphic and acquires, at the same time, upper-jaw characters such as the presence of vibrissae (Ozeki, et al., 2004; Clouthier, et al., 1998)’(Ruest, et al., 2004; Yanagisawa, et al., 2003; Charite, et al., 2001). In *Edn1*, *Ednra* or *Hand2* mutants the lower jaw is transformed, but a dorso-ventral symmetry of the mouth is not acquired. Conversely, activation of the Edn-signalling pathway in the PA1 maxillary NCCs contingent, where it is normally silent, results in the activation of *Dlx5/6* and *Hand2* leading to the transformation of upper jaws into dentary-like structures articulated with unaffected lower jaws presenting their own Meckelian cartilages (Sato, et al., 2008b). Importantly, it has been shown that Dlx5/6-positive NCCs do not only contribute to the morphogenesis of bones and cartilages, but exert also an instructive role for the patterning and differentiation of the myogenic mesoderm; the constitutive inactivation of *Dlx5/6* results in absence of masticatory muscles while in *Ednra* mutants, the Edn1-independent reactivation of *Dlx5/6* in PA1 is sufficient to rescue muscular differentiation (Heude, et al., 2010). The sum of these results suggests that *Dlx5/6* expression in different jaw precursors including, *ad minima*, NCCs and distal ectodermal cells, has a critical role in sculpturing vertebrate facial structures. Here, by genetically down- or up-modulating the expression of *Dlx5/6* specifically in NCCs, we addressed the question of which morphological messages are conveyed by NCC-restricted *Dlx5/6* expression and which other cues depend from their expression in other cell types.

We show that, if *Dlx5/6* are expressed in either mandibular or maxillary NCCs the morphological and molecular signature of a lower jaw can be recognized while, in their absence, the jaws assume invariably a maxillary identity. In particular the main morphological elements which we have considered as typical traits of mandibular identity are: 1) the presence of a Meckelian cartilaginous rod, 2) the absence of midline fusion between the right and left jaws that are joined only at the distal symphysis, 3) the absence of an infraorbital foramen, 4) the presence of an elongated incisor, 5) the absence of a premaxillary bone, 6) the absence of vibrissae, 7) the presence of tongue and associated musculature and 8) the insertion of masticatory muscles. Maxillary identity on the contrary has been defined by the following characters: 1) the absence of a Meckelian cartilage, 2) the fusion of dermocranial bones along the midline to form a palate which might present palatine rugae, 3) the presence of an infraorbital foramen, 4) lateral extensions of the bones forming zygomatic-like structures, 5) the presence of a premaxillary bone, 6) the presence of vibrissae aligned in five regular lines, 7) short incisors supported by the premaxillary bone and not by other maxillary components, 8) the connection of the superficial masseter to the infraorbital foramen.

Remarkably, all mandibular traits are invariably present when *Dlx5/6* are expressed in NCCs either in the upper or in the lower jaw. Conversely, maxillary traits appear in the absence of *Dlx5/6* expression in NCCs associated to a defective muscular system.

Our first conclusion is that NCC-limited expression of *Dlx5/6* is necessary and sufficient to specify a mandibular identity and to maintain the myogenic program for the formation of muscularized jaws.

The second, flagrant, result of this study is that *Dlx5/6* must be expressed in cellular contingents different from NCCs to permit the development of matching, functional jaws. Indeed, the inactivation of *Dlx5/6* only in NCCs transforms the lower jaw into an hypomorphic upper jaw that is severely out of register from the existing maxilla. On the contrary, inactivation of *Dlx5/6* in all cell types, including NCCs results in the transformation of both upper and lower jaws into similar matching structures that, if muscularized, could prefigure a functional mouth. It must be pointed out that transient expression of the transcription factors *Dlx5/6* might be sufficient to change the developmental trajectory a cellular contingent and might confer the instructive capacity needed to modulate morphogenesis. In line with this concept, we have previously shown that the transient expression of *Dlx5/6* in epithelial precursors deriving from the lamboid junction (the intersection between presumptive maxillary and frontonasal bud-derived structures) is needed for correct morphogenesis of the upper jaw (Gitton, et al., 2014). The main findings of this paper are summarized in Fig. 9.

**Figure 9:**
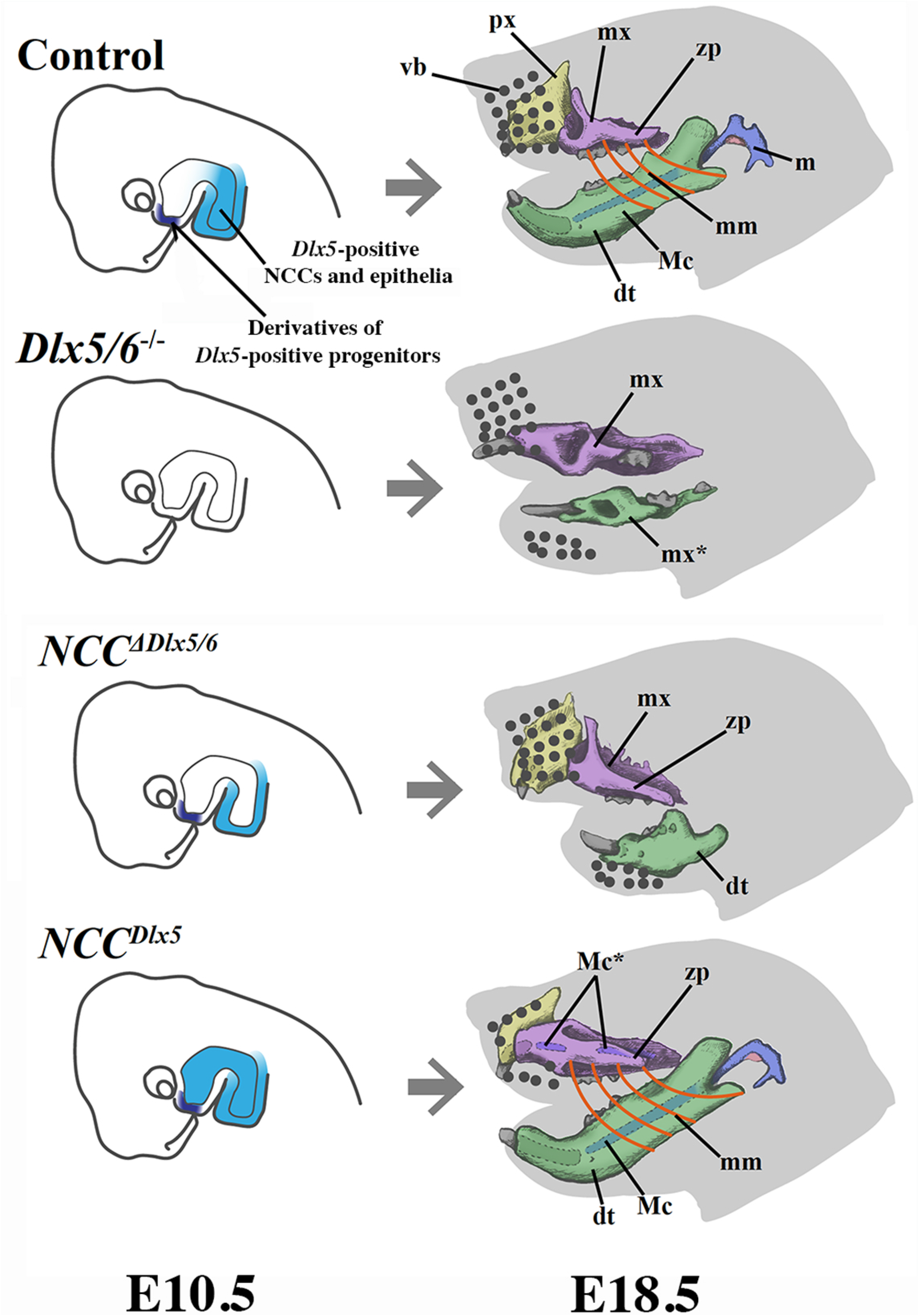
Summary diagram of the molecular (E10.5) and morphological (E18.5) alterations obtained after inactivation or induction of *Dlx5/6* expression in NCCs. Abbreviations: dt, dentary bone; m, malleus; Mc, Meckel’s cartilage; mm, masseter muscle; mx, maxillary bones; px, premaxilla; vb, vibrissae; zp, zygomatic process. * transformed ectopic elements.

It has been shown that FGF8 signals deriving from the mandibular PA1 epithelium are essential for jaw morphogenesis. Indeed, inactivation of mouse *Fgf8* in PA1 epithelia results in the loss of most PA1 skeletal derivatives (Trumpp, et al., 1999). Interestingly, mandibular, but not maxillary ectomesenchyme is competent to respond to Fgf8 maintaining *Dlx5* expression (Ferguson, et al., 2000) (Park, et al., 2004). A possible interpretation of our data would be that *Dlx5* expression in the lower jaw epithelium could be essential to maintain molecular signals such as Fgf8 and, in so doing, could permit correct lower jaw morphogenesis. Interestingly, at the end of gastrulation *Dlx5* expression first appears in lateral bands of the embryonic/extraembryonic junction at the late-streak stage which precedes neural plate formation (Yang, et al., 1998). Later, at E7.25 expression of *Dlx5* is detected in the rostral and in the posterior ectoderm. At even later stages, *Dlx5* is found in NCC progenitors and in epithelial precursors of the ventral cephalic ectoderm (VCE). During migration, NCCs down-regulate *Dlx5* expression and reactivate it only after arrival in the mandibular PA1 in response to an Edn1 signal. At E8.5 the NCCs-derived mesenchyme of the early PA1 is clearly Dlx5-positive (Acampora, et al., 1999), however the epithelial component of PA1 does not express yet express *Dlx5* (Yang, et al., 1998). The lower jaw epithelium seems to activate *Dlx5* expression between E8.5, when the epithelium is still Dlx5-negative (Yang, et al., 1998), and E10.5, when the epithelial *Dlx5* expression is comparable to that of the ectomesenchyme (Vincentz, et al., 2016), (see also Fig. 9). It seems plausible therefore that *Dlx5* expression both in NCCs and in the PA1 mandibular epithelium is essential to permit correct lower jaw morphogenesis. VCE cells also down regulate *Dlx5* expression, but their derivatives migrate then toward anterior parts of PA1 (Gitton, et al., 2014) and are essential for correct maxillary development. It seems therefore that the integration of signals deriving from three *Dlx5*-positive cellular progenitors (NCCs, PA1 epithelial cells and VCE cells) is needed for the generation of functional, matching jaws. *Dlx5* expression in these different territories might well depend on different regulatory mechanisms. An example of this differential regulation is the observation that the *I56i* enhancer is active in PA1 NCCs, but not in epithelial structures such as the VCE, the PA1 epithelium or the otic vesicle (Park, et al., 2004).

Vital biological functions such as feeding or reproduction often depend on the interaction between matching structures located either in the same individual (e.g. jaws) or in separate members of the same species (e.g. sexual organs). The coevolution of synergistic complementary structures in separate individuals or in different territories of the same organism has, therefore, been instrumental to ensure species survival. We show that, for the specific case of mouth development, the same cluster of genes is: 1) determining lower jaw identity and 2) tuning the morphogenetic process of both the upper and the lower jaw generating a mouth endowed with opposing and matching mandibles capable to support feeding and/or predation. These two levels of morphogenetic regulation must be coordinated to generate a functional oral organ capable of mastication. These morphogenetic processes occur in different cell types located in spatiotemporally separate embryonic territories suggesting that diverse *Dlx5/6*-depending pathways are involved. At least in the case of the jaws, the co-evolution of synergistic anatomical structures ostensibly involves the coordination of different regulations involving the same genes in distant embryonic territories.

These observations open very profound questions on how these different regulations might have evolved and be selected. It is remarkable however that both NCCs and the VCE derive from the same *Dlx5*-positive early progenitors at the very end of gastrulation suggesting interesting evolutionary scenarios that still need to be understood. Probing the origin of matching, muscularized, jaws could open a window to understand more generally how different harmonious, but separate, parts of the body coordinate their morphogenesis.

## MATERIALS AND METHODS

### Mice

Procedures involving animals were conducted in accordance with the directives of the European Community (council directive 86/609), the French Agriculture Ministry (council directive 87–848, 19 October 1987, permissions 00782 to GL) and approved by the University of Tokyo Animal Care and Use Committee and by the “Cuvier” ethical committee (approval n° 68-028r1). Mice were housed in light, temperature (21°C) and humidity controlled conditions; food and water were available ad libitum. WT animals were from Charles River France. Double *Dlx5* and *Dlx6* (*Dlx5/6*) null mice (Beverdam, et al., 2002; Merlo, et al., 2000) and the double conditional mutant *Dlx5/6^flox/flox^* (Bellessort, et al., 2016) in which the DNA-binding region of both *Dlx5* and *Dlx6* is deleted by cre-recombinase were maintained and genotyped as reported. The inducible Cre driver strain *Wnt1-creERT2* (Stock #008851) and *Wnt1-cre* (Stock #022501) were purchased from Jackson Laboratory Maine, USA through Charles River Laboratories (L’Arbresle, France) and maintained on a C57BL/6J genetic background.

To obtain mice carrying the *R26R^CAG-flox-DLX5/+^* allele, an *F3/*FRT-flanked cassette containing the CAG promoter, a floxed stop sequence, Flag-tagged mouse *Dlx5* cDNA and a poly(A) additional signal were inserted into the targeting vector pROSA26-1 (P. Soriano, Mount Sinai School of Medicine, New York, NY, USA) (Addgene, plasmid 21714). Homologous recombination was performed on the ROSA26 locus of B6129F1-derived ES cells. Targeted ES clones were injected into ICR blastocysts to generate chimeras. Chimeras were crossbred with ICR females. *R26R^CAG-flox-DLX5/+^* mice were crossed with *Wnt1-cre* mice (Chai, et al., 2000) to induce NCC-specific expression of *Dlx5*(*NCC^Dlx5^* mice).

Embryos were obtained from pregnant dams, washed and dissected in ice-cold phosphate buffered saline (pH7.4). Embryos selected for fixation were immersed in 4% paraformaldehyde from 15mn to 2h, photographed and further processed (Bellessort, et al., 2016; Sato, et al., 2008b). Specimens selected for biochemical analyses were dissected in PBS and processed for RT-qPCR, Affymetrix, Western blot and Southern blot as described in Supplementary text 1 and (Bellessort, et al., 2016).

Intraperitoneal tamoxifen (Sigma-Aldrich, France, #T5602) injections were performed as previously described (Gitton, et al., 2014) using 3 to 5mg/day/pregnant mouse. To cover most of the period of neural crest delamination and migration (Danielian, et al., 1998) *Dlx5/6^Fl/Fl^* ::*Wnt1^Cre-ERT2/+^* pregnant dams received two IP injections of tamoxifen at E6 and E7, generating *NCC*^Δ*Dlx5/6*^ embryos.

### Histology and 3D reconstruction

Heads from E18.5 (mutant and wild type foetuses were fixed in Bouin’s solution (Sigma, France), embedded in paraffin and complete sets of frontal or parasagittal serial sections (12μm) were prepared. All sections were stained by Mallory’s trichrome (Heude, et al., 2010) and photographed (Nikon Digital Site DS-FI1). Pictures were aligned, piled and registered using the Fiji plug-in of NIH ImageJ “Register Virtual Stack Slices” (http://fiji.sc/wiki/index.php/Register_Virtual_Stack_Slices). 3D segmentation was performed with Mimics (Materialise, Belgium: http://biomedical.materialise.com/mimics) and visualized using Adobe Acrobat 9 pro.

### Skeletal preparation and staining

Alizarin red/ alcian blue staining was performed, as previously described (McLeod, 1980). Samples were fixed in 95% ethanol for a week, permuted to acetone for three days and incubated with 0.015% alcian blue 8GS, 0.005% alizarin red S and 5% acetic acid in 70% ethanol for three days. After washing with distilled water, the samples were cleared in 1% KOH for several days and in 1% KOH glycerol series until the surrounding tissues turned transparent. The preparations were stored in glycerol.

### *In situ* hybridization

Whole-mount in situ hybridization was performed, as described previously (Heude, et al., 2010). Embryos were fixed one overnight in 4% paraformaldehyde in PBT. After dehydration and rehydration with methanol, the embryos were bleached for 1 hour in 7.5% H2O2 in PBT and then washed in PBT 3 times. The samples were treated with 5mg/ml proteinase K for 40 seconds at room temperature, treated with 2mg/ml glycine in PBT to stop the enzyme reaction, and post-fixed in 4% paraformaldehyde and 0.2% glutaraldehyde for 20 minutes on ice. After the pretreatment, the samples were pre-hybridized for more than 1 hour at 70°C in hybridization mix (50% formamide, 5 × SSC (1 × SSC is 0.15 M NaCl plus 0.015 M sodium citrate), 1% SDS), 50 mg/ml heparin and 50 mg/ml yeast tRNA. With digoxygenin-labeled RNA probe in hybridization mix, the samples were hybridized overnight at 70°C. The samples were then washed 3 times in hybridization mix at 70°C, then in 0.2 M NaCl, 10 mM Tris-HCl (pH7.5), 0.1% Tween-20 for 5 minutes ant treated with 100 mg/ml RNase for 30 minutes at 37°C. After a final wash in 50% formamide, 2 × SSC for 1 hour at 65°C, the samples were pre-blocked with sheep serum, incubated with alkaline phosphatase-conjugated anti-digoxygenin antibody, and stained with nitro blue tetrazolium and 5-bromo-4-chloro-3-indoyl phosphate. Probes were prepared by RT-PCR and used in published (Kitazawa, et al., 2015; Sato, et al., 2008b).

### Quantitative real-time RT-PCR

The maxillary and mandibular processes were dissected from E10.5 control and *NCC^Dlx5^* mice. Total RNA was extracted from five sets of PAs by ISOGEN-II (Nippon Gene). One-μg samples were then reverse-transcribed using ReverTra Ace (TOYOBO) with RS19-15dT primer. Quantification of amount of each mRNA was performed by real-time PCR analysis using a LightCycler (Roche) and Real-Time PCR Premix with SYBR Green (RBC Bioscience) following the manufacturer’s protocol. Glyceraldehyde-3-phosphate dehydrogenase (*Gapdh*) was used as internal control. PCR was performed using following primers, Dlx5 previously described (Vieux-Rochas, et al., 2010), 5’-AGACAGCCGCATCTTCTTGT-3’ and 5’-CTTGCCGTGGGTAGAGTCAT-3’ for *Gapdh*.

### Gene expression profiling

The maxillary process, the mandibular process and the PA2 were collected from E10.5 control and *NCC^Dlx5^* mice and subjected to Affymetrix GeneChip analysis. Each sample was a mixture from 3 littermates. Preparation of the cRNA and hybridization of probe array were performed on an Affymetrix GeneChip Mouse 430 2.0 array which contains 45,101 probe sets according to the manufacturer’s instructions (Affymetrix, Santa Clara, CA). The expression value for each mRNA was obtained by the Robust Multi-array Average (RMA) method. The gene set probes were filtered on an expression (20.0–100.0) percentile. Genes with the expression level lower than 20.0 percentile at least in one sample were eliminated from the analysis. After excluding the probes whose gene symbols were not identified, about 35,000 genes remained and used for further analysis. Annotation of the probe numbers and targeted sequences are shown on the Affymetrix web site.

## ACKNOWLEDGEMENTS

This research was partially supported by the EU Consortium HUMAN (EU-FP7-HEALTH-602757) and by the ANR grants TARGETBONE (ANR-17-CE14-0024) and METACOGNITION (ANR-17-CE37-0007) to GL, and an ATM grant (FORMs) to YG, CdL is supported by a grant of the French Ministry of Research. This work was supported in part by grants-in-aid for scientific research from the Ministry of Education, Culture, Sports, Science and Technology, Japan (15H02536) and Core Research for Evolutional Science and Technology (CREST) of the Japan Science and Technology Agency (JST), Japan (JPMJCR13W2).

A particular thank goes to the team in charge of mouse animal care and in particular M Stéphane Sosinsky and M Fabien Uridat and to Pr. Amaury de Luze in charge of animal wellbeing. We thank the Stanford Exchange Program Scholars Mss. Shaheen Jeawoody, Andrea Fontaines and Heydi Malavé for the excellent help given to this work and Mss. Aicha Bennana and Lanto Courcelaud for administrative assistance.

## SUPPLEMENTARY FIGURES LEGENDS

**Figs. Supplementary 1_4: 3D pdf files to be opened in Acrobat, which allows selection and manipulation of 3D reconstructed craniofacial structures described in this study.**

Colour code as in Figs. 3 and 4: Yellow, premaxillary bone; purple, maxillary bones; green, dentary bone; blue Meckelian cartilage; red, tongue; grey, teeth. Using the Acrobat functions different structures can be visualized or not, or shown with different options.

**Fig. S1:** Control craniofacial skeleton at E18.5

**Fig. S2:** *Dlx5/6^-/-^* craniofacial skeleton at E18.5

**Fig. S3:** *NCC*^Δ*Dlx5/6*^ craniofacial skeleton at E18.5

**Fig. S4:** *NCC^Dlx5^* craniofacial skeleton at E18.5

**Fig. S5:**
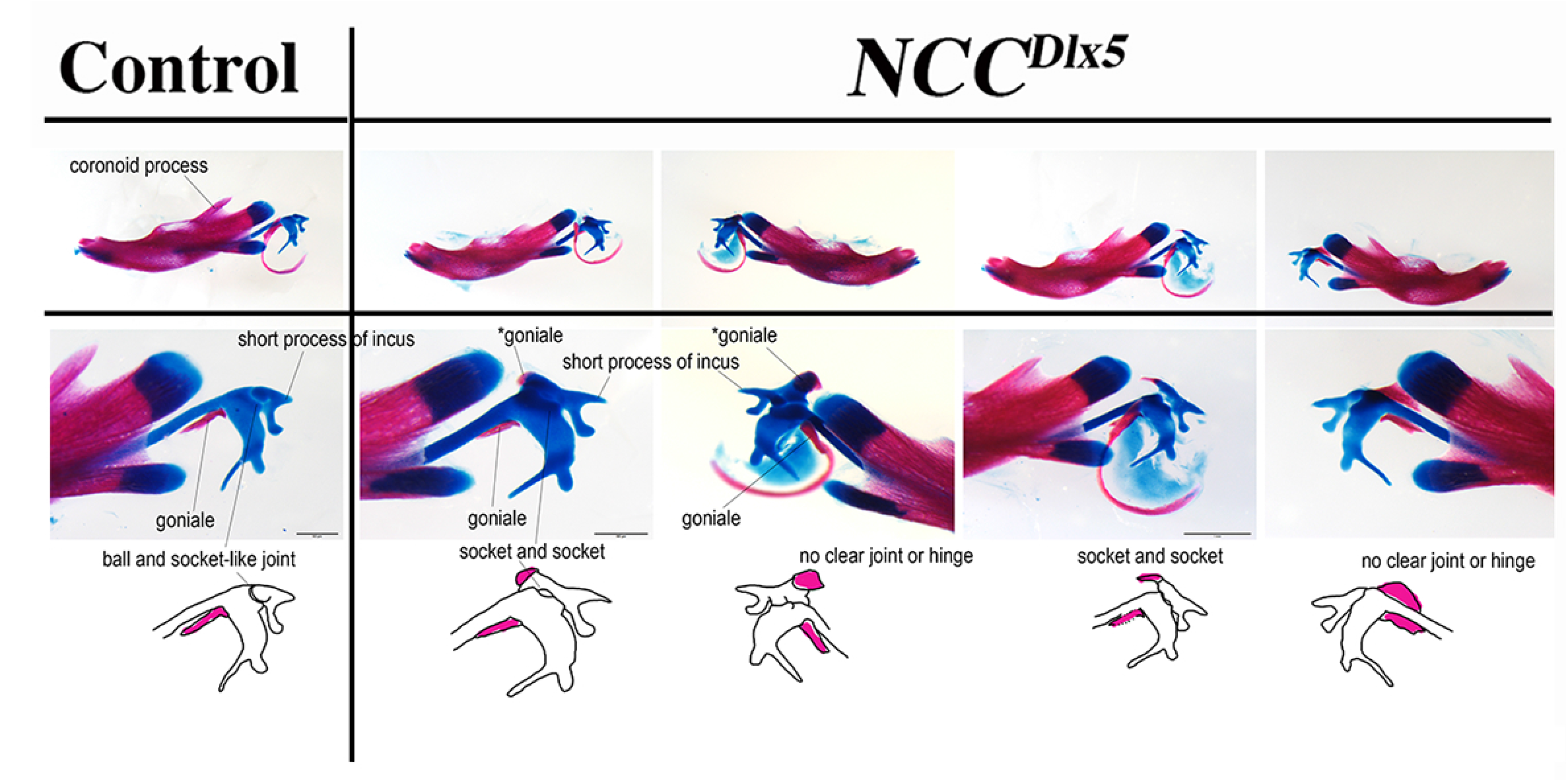
Morphological analysis of the transformation of the incus/malleus region in *NCC^Dlx5^* E18.5 foetuses. The elongation of the short process of the incus, the presence of a small ectopic bone, which could be interpreted as a duplicated gonial bone, adjacent to the incus, and the fact that the malleus-incus joint, which is normally of ball-and-socket type, is symmetric suggest a partial transformation of the incus in a malleus-like structure.

